# The AMD-associated genetic polymorphism CFH Y402H confers vulnerability to Hydroquinone-induced stress in iPSC-RPE cells

**DOI:** 10.1101/2024.11.06.622214

**Authors:** Angela Armento, Inga Sonntag, Ana-Cristina Almansa-Garcia, Merve Sen, Sylvia Bolz, Blanca Arango-Gonzalez, Ellen Kilger, Ruchi Sharma, Kapil Bharti, Rosario Fernandez-Godino, Berta de la Cerda, Simon J Clark, Marius Ueffing

## Abstract

Age-related macular degeneration (AMD), a degenerative disease of the macula, is caused by an interplay of diverse risk factors (genetic predisposition, age and lifestyle habits). One of the main genetic risks includes the Y402H polymorphism in complement Factor H (FH), an inhibitor of complement system activation. There has been, and continues to be, much discussion around the functional consequences of this Y402H polymorphism, whether the soluble FH protein confers its risk association, or if the cells expressing the protein themselves are affected by the genetic alteration. In our study, we examined the cell characteristics of the retinal pigment epithelium (RPE) cells, which play a major role in retinal homeostasis and stability and which are synonymously linked to AMD.

Here, we employ RPE cells derived from induced pluripotent stem cells (iPSC) generated from donors, carrying either homozygous 402Y (low risk) or 402H (high risk) variants of the *CFH* gene. RPE cells were treated with Hydroquinone (HQ), a component of cigarette smoke, to induce oxidative damage. Intriguingly, RPE cells carrying high genetic risk proved more vulnerable to oxidative insult when exposed to HQ, as demonstrated by increased cytotoxicity, caspase activation, compared to the low-risk RPE cells.

The exposure of RPE cells to RPE conditioned medium, normal human serum (NHS) and inactivated NHS (iNHS) had minimal impact on cell cytotoxicity and caspase activation, nor did the presence of purified soluble FH rescue the observed effects. This suggests that the degree of cellular susceptibility to oxidative stress is not conferred by soluble FH protein and other complement sources, but intercellularly because of the corresponding genetic risk predisposition. Considering the known connection of oxidative stress to proteotoxic stress and degrading processes, we investigated the unfolded protein response (UPR) and autophagy. When exposed to HQ, RPE cells showed an increase in autophagy markers, however iPSC-RPE cells carrying high genetic risk showed an overall reduced autophagic flux. Our data support the hypothesis that RPE cells carrying high genetic risk are less resilient to oxidative stress.

## Introduction

Age-related macular degeneration (AMD) is a complex and progressive disease of the macula, mainly affecting the elderly population, culminating in vision loss. As the number of cases is expected to increase, threatening the patient’s quality of life, AMD places a major burden on national health systems [1]. From a clinical perspective, late-stage AMD is divided into wet AMD or choroidal neovascularization (CNV) and dry AMD, which accounts for the majority of AMD cases and for which therapeutic options are limited. Dry AMD is defined by geographic atrophy (GA), characterized by degeneration of the photoreceptors (PR) and underlying Retinal Pigment Epithelium (RPE) cells [1]. RPE cells hold several functions crucial to maintaining retina visual function and to supporting PR function. As a cellular monolayer interfacing the neuroretina and the choroidal blood supply space, RPE cells are responsible for nutrient supply, fluids, and discard material exchange, including the processing of shed PR outer segments (POS). Moreover, RPE cells possess a variety of antioxidant mechanisms to protect the retina from excessive oxidative stress [2]. The exact mechanism by which RPE cells become dysfunctional in AMD is not fully understood and most importantly how the combination of AMD risk factors interacts to contribute to RPE disease phenotypes remains quite complex to understand. Besides aging, which is the primary cause of AMD, unhealthy life-style choices, such as smoking, contribute to disease progression. Moreover, genetic predisposition plays a crucial role in AMD risk [3].

Genome-wide association studies (GWAS) have identified several associations between genetic variance and AMD risk and progression. Several risk loci have been identified as modifying risk for disease, but two main risk loci predominate: one located in chromosome 10 (Chr10q26) around the *ARMS2/HTRA1* genes [4], and the other in chromosome 1 (Chr1q32) known as the Regulators of Complement Activation (RCA) cluster: that contains genes regulating the complement system [5]. This RCA cluster includes the Complement Factor H (*CFH*) gene, encoding for the Factor H (FH) protein and its truncated splicing variant FHL-1 [6]. A very common polymorphism, leading to a Y402H amino acid change in the FH/FHL-1 proteins, is a defining feature of the genetic haplotype with the strongest association to AMD risk.

The FH/FHL-1 proteins function as negative regulators of the complement system, by binding to host surfaces and helping deactivate the central protagonist of complement activation, C3b [7]. A proteolytic enzyme, complement factor I (FI), can bind and cleave C3b, inactivating it to produce iC3b. However, FI can only achieve this in the presence of a co-factor. Despite several different cell-bound FI co-factors, FH and FHL-1 represent the only soluble FI co-factors that can provide protection to a host’s acellular surfaces, such as basement membranes and the glycocalyx [8]. The Y402H polymorphism occurs within a major anchoring domain of FH and FHL-1, and is believed to hamper the proteinś ability to bind surfaces such as Bruch’s membrane [9]. Indeed, markers of complement over-activation are seen in the eyes of human donors carrying genetic risk haplotypes including the Y402H polymorphism [10]. Interestingly, markers of complement over-activation can also be found systemically in patients with AMD, although some debate remains around whether this is a systemic phenomenon, or a local, tissue-specific, event that can be detected systemically [11].

However, in recent years a novel concept of non-canonical FH functions has been introduced [12]. This includes a number of cellular functions which have been suggested to be regulated by FH, mainly in its intracellular form and most likely independent from its function as complement system inhibitor in the extracellular space [12, 13]. Recent studies have highlighted that FH, and the Y402H polymorphism, could contribute directly to RPE pathology. It was shown that iPSC-RPE carrying the Y402H polymorphism presents mitochondria metabolism impairments [14], mitochondria abnormalities [15], and an accumulation of lysosomes and phagosomes [15, 16]. As RPE cells and neuroretina coexist in a metabolic balance with each other, these alterations could affect the ability of RPE cells to metabolically support the retina and process the discard material deriving from the PRs *via* the autophagy-lysosomal flux [17, 18]. No less importantly, these altered features could render the RPE cells even more dysfunctional in AMD-like conditions, such as increased oxidative stress. Indeed, several factors contributing to AMD often culminate in excessive oxidative stress, as a consequence of aging or cigarette smoke exposure [19]. Accumulation of Reactive Oxygen Species (ROS) can damage cellular homeostasis, leading to the buildup of damaged organelles and proteins by oxidation [20, 21]. Healthy RPE is quite resistant to the oxidative damage throughout its lifetime, and can continue to operate degrading pathways, such as autophagy and endoplasmatic reticulum-associated degradation (ERAD), to process and expel the damaged material [22]. However, RPE cells that present impaired mitochondria, such as iPSC-RPE cells carrying the Y402H polymorphism of FH [15], may be predisposed to a less efficient oxidative stress response and therefore may not be able to survive an excessive oxidative stress environment. In these circumstances, RPE dysfunction will be elevated and pave the way to PR degeneration, ultimately being the cause of vision loss.

In accordance with this hypothesis, in our previous work, we have shown that hTERT-RPE1 cells deprived of FH via silencing, are more vulnerable to oxidative stress insult (H_2_O_2_), involving alteration in metabolic pathways, as oxidative phosphorylation and glycolysis, mitophagy and inflammation [23].

To move forward and better understand the influence of FH dysregulation in RPE pathology during AMD progression, we employed, in this work, iPSC-RPE cells derived from either low or high genetic risk donors, which closely represent the maturity of RPE cells *in vivo*. To investigate the impact of oxidative stress in mature RPE cells, we employed a more physiological oxidative agent: Hydroquinone (HQ). HQ is a benzene metabolite that has high redox activity and is a well-known substance used to induce oxidative stress [24]. HQ is a component of cigarette smoke and indeed elevated plasma levels of HQ have been found in smokers compared to non-smokers [25]. In AMD research, both *in vitro* and *in vivo* studies proved that exposure to HQ causes oxidative damage and AMD-like pathology in mice and immortalized RPE cell lines [26, 27].

In this work, we investigated the differential stress response of human iPSC-RPE cells to oxidative stress conferred by Hydroquinone. We analyzed differentiated human RPE, where low-risk (LR) cells are homozygous carriers of the FH 402Y variant, while high-risk cells (HR) carry the FH 402H variant. We demonstrate that RPE cells carrying the FH 402H variant are more vulnerable to HQ-induced oxidative stress in a concentration-dependent manner, affecting autophagy as well as increasing apoptosis independent of extracellular levels of complement.

## Methods

### Induced pluripotent stem cells and RPE differentiation

Human induced pluripotent stem cells (iPSC) were obtained from different sources (specified in Suppl. Figure 1A). iPSC cell lines 1, 2, 4, 5, and 6 were obtained from genetically screened human donors as explained previously [16]. iPSC cell lines 3, 7, 8 were generated as previously described [28] and purchased from the Spanish National Stem Cell Bank. iPSCs were grown on hESC-matrigel (354277, Corning) coated plates in E8 medium, medium was changed daily. iPSCs were differentiated to RPE cells according to previously described protocol [29, 30]. Briefly, iPSCs were seeded on hESC matrigel (356237, Corning) in differentiation medium (DMEM/F12, 31331, Gibco; N2 supplement, 17502, Gibco; B27 supplement, 17504, Gibco; NEAA, 11140, Gibco; Knock Out SR, 10828, Gibco). Briefly, medium was replaced every two days for 14 days following specific formulations: Nicotinammide 10 mM (Day 0-4, NO636-100G, Sigma-Aldrich), Noggin 50ng/ml (day 0-4, 1967-NG-025/CF, R&D systems), DKK-1 10 ng/ml (day 0-6, 5439-DK-010/CF, R&D Systems), IGF-1 10ng/ml (day 0-6, 1291-G1-200, R&D Systems), bFGF 5 ng/ml (day 2-4, AF-100-18B, Peprotech), activinA 100 ng/ml (day 4-14, 120-14E, Peprotech), SU5402 10 μM (day 6-14, sc-204308, Santa Cruz).

At day 14, cells were enriched in medium containing rock inhibitor (StemMACS Y27632), and seeded in Geltrex (A1413302, Gibco) or hESC-matrigel coated plates for expansion. Medium was changed twice a week and RPE cells were maturated for at least 30 days. RPE cells were passaged using TrypLE protocol according to manufacturer instructions (12604; Thermo Fisher Scientific). Cells were used for experiments from passage 2 to passage 4 in the desired plate format (12 well-culture inserts, 96-well plates, 12-well plates) and were cultured for at least 50 days before experiments. Cell culture treatments were carried for 48hours before analyses and included: Hydroquinone (HQ) 100-200 μM (H9003, Sigma-Aldrich), Tunicamycin 5μg/ml (SML1287, Sigma-Aldrich), Bafilomycin (50 nM, B1793, Sigma), purified FH 1μg/ml (A137, Comptech), Normal human serum 1% (H4522, Sigma-Aldrich), inactivated human serum 1% (H3667, Sigma-Aldrich).

### Western Blot

Cellular lysates were collected in IP lysis buffer (87787, Thermo Fisher Scientific) with protease and phosphatase inhibitors (1861281, Thermo Fisher Scientific). After 30 minutes incubation at 4°C, followed by centrifugation, Protein concentration was measured according to the Bradford method following manufacturer instructions (500-0006, Biorad). Proteins were resuspended in NuPAGE™ LDS Sample Buffer containing reducing agent (NP0007, Invitrogen, California, USA), separated on 8-16% SDS-PAGE gels (XP08165Box; Thermo Fisher Scientific) and transferred on PVFD membranes (0,45µm, 10600023, Cytiva). Membranes were exposed overnight to the primary antibodies (anti-FH, sc-166608, Santa Cruz, Texas, USA; anti-Bip, #3177, Cell Signalling; anti-IRE1a, #3294, Cell Signalling; anti-LC3, 0231-100/LC3-5F10, Nano tools; anti-SQSTM1, BML-PW9860-100, Enzo; anti-Bax, #12105, Cell Signalling; anti-Bcl-2, #2870, Cell Signalling; anti-Sod2, #13141, Cell Signalling; anti-Tubulin, #2148, Cell Signalling) and for 1 hour to HRP-conjugated anti-mouse or anti-rabbit secondary antibody (1:2.000, 7076S, 7074P2, Cell Signaling, Massachusetts, USA). Immunoreactivity was visualized with Pierce™ ECL Western Blotting Substrate (32106, Thermo Fisher Scientific, Massachusetts, USA) and detected with FusionFX instrument (Vilber Lourmat, France).

### RNA extraction, cDNA synthesis and qRT-PCR

Cell pellets resolved in Purezol (732-6880, Biorad, Germany), homogenized by inversion, and incubated at room temperature for 5 minutes. After addition of chloroform, samples were vortexed for 15 seconds, incubated 5 minutes at room temperature, and then centrifuged at 12,000 g for 15 minutes at 4°C. The aqueous phase was collected and mixed with isopropanol for precipitation. Samples were centrifuged at 12,000 g for 15 minutes at 4°C. Pellets were rinsed twice with EtOH 75%, dried, resuspended in 20 μl of RNase-free water. RNA purity and concentration were measured using Nanodrop. cDNA was synthesized via reverse-transcription of 2-5 μg of RNA using M-MLV Reverse Transcriptase (200 U, M1705, Promega, Wisconsin, USA), random primers (10 ng/μl, C1181, Promega, Wisconsin, USA) and dNTPs (0.5 mM, U1515, Promega) in a total volume of 20 μl. cDNA was used to analyse differences in gene expression by qRT-PCR employing iTaq Universal SYBR Green Supermix (1725122, Biorad, Germany) along with gene-specific forward and reverse primers (250 nM) according to manufacturer instructions. Relative mRNA expression of each gene of interest (GOI) was quantified by using PRPL0 as the housekeeping control gene. Primers are listed below: PINK1 (fwd 5’-GGC TTG GCA AAT GGA AGA AC −3, rev 5’-CTC AGT CCA GCC TCA TCT ACT A −3), PARKIN (fwd 5’-CCA CAC TAC GCA GAA GAG AAA −3’, rev 5’-GAG ACT CAT GCC CTC AGT TAT G −3’), PPARGC1A (fwd 5’-AGA GCG CCG TGT GAT TTA T −3’, rev 5’-CTC CAT CAT CCC GCA GAT TTA −3’), CAT (fwd 5’-CTG GAG CAC AGC ATC CAA TA −3’, rev 5’-TCA TTC AGC ACG TTC ACA TAG A −3’), GPX1 (fwd 5’-CAT CAG GAG AAC GCC AAG AA −3’, rev 5’-GCA CTT CTC GAA GAG CAT GA −3’), CFH (fwd 5’-CTG ATC GCA AGA AAG ACC AGT A −3’, rev 5’-TGG TAG CAC TGA ACG GAA TTA G −3’), PRPLO (fwd 5’-GGA GAA ACT GCT GCC TCA TAT C −3’, rev 5’-CAG CAG CTG GCA CCT TAT T −3’).

### ELISA

Cell culture media from the apical and basal compartments of RPE culture inserts were collected. Cellular debris were removed by centrifugation. PEDF ELISA (RD191114200R, BioVendor) was performed according to manufacturer instructions. Freshly prepared standards, quality controls, and samples were incubated in ELISA plates at room temperature on shaker for 1 hour. After washes, Biotin-labelled antibody was added to each well and incubated at room temperature on shaker for 1 hour. Then, following washes, the Streptavidin-HRP Conjugate was added into each well and incubated as before. Signal reaction was initiated by adding the detection substrate and stopped after 5 minutes with Stop Solution. Absorbance was measured at 405 nm at the Spark multimode microplate reader (Tecan, Switzerland).

### TER

TER measurements are performed with a Millicell-ERS-2 meter according to the manufacturer’s instructions. The electrodes were applied on the apical and basal sides of iPSC-RPE, the current was passed, and resistance values (Ohms) were noted down for each well. The average of three measures was recorded for each well. Actual resistance (Ohms*cm^2^) is calculated by multiplying TER to the area of the insert.

### Immunostaining

iPSC-RPE monolayers grown on transwells were fixed in 4% PFA at room temperature. Small sections were selected and used for immunostaining. Briefly, samples were washed in PBS for 10 minutes and incubated 5 minutes with 0.3% Triton. After two rinses in PBS, samples were shortly incubated with TrueBalck solution (23007, Biotium). After three rinses in PBS, samples were blocked in NGS (normal goat serum, S26, EMD Millipore Corp., USA) for 1 hour and then exposed overnight at 4°C to the primary antibody (ZO-1, 610966, BD transduction). Following three washes in PBS, samples were incubated with secondary antibody (Goat anti-mouse Alexa 568, A11031, molecular probes). Samples were mounted on coverslips with Fluoromount-G solution (17984-25, Electron Microscopy Sciences, Pennsylvania, USA). Images were obtained using Z-stacks on a Zeiss Axio Imager Z1 ApoTome Microscope.

### In vitro assays

Cytotoxicity and caspase activity were assessed using the MultiTox-Fluor Assay (G9201, Promega) and the Caspase-Glo Assay (G8091, Promega). Cytotoxicity, defined by cell membrane damage, was assessed by cell-impermeable bis-AAF-R110 (bis-alanylalanyl-phenylalanyl-rhodamine 110) dye, which is cleaved by dead-cell proteases released in the cell culture supernatants after membranes damages. Fluorescence is read at 485Ex/520Em. After detection, caspase 3/7 reagent was added in the wells and incubated for 10 minutes at room temperature in the dark. Equal volumes were transferred to OptiPlate-96 (6005290, Perkin Elmer) and luminescence signals were measured. Spark multimode microplate reader (Tecan, Switzerland) was used for fluorescence and luminescence measurements.

Levels of H_2_O_2_ were measure using the ROS-Glo H_2_O_2_ assay (G8820, Promega) and assay was performed according to manufacturer instructions. Briefly, cells were exposed to the H_2_O_2_ substrate and incubated in the incubator for 4 hours. At this point, H_2_O_2_detection buffer (containing cysteine and signal enhancer) was added, incubated 10 minutes at room temperature in the dark under shaking. Equal volumes were transferred to OptiPlate-96 and luminescence was measured.

CellTiter-Glo assay (G9241, Promega) was used to measure total ATP levels present in the cultures. CellTiter-Glo reagent was added to the cells, incubated at room temperature for 10 minutes. Equal volume were transferred to OptiPlate-96 and luminescence was measured. JC1 dye (T3168, ThermoFisher Scientific) was used to monitor mitochondrial health, as exhibits differential accumulation depending on mitochondria membrane potential. At lower potential JC-1 is present as monomer and it is indicated by green fluorescence, while at higher membrane potential JC-1 aggregates in the mitochondria and emits in the red fluorescence. Therefore, a JC-1 red/green ratio is indicative of mitochondria polarization. The cells were exposed to the JC-1 dye for 15 minutes in the incubator and rinsed three times in PBS and fluorescence signal were measured in the RPE layer at the Spark multimode microplate reader (Tecan, Switzerland).

### Transmission Electron Microscopy

iPSC-PRE samples were fixed in 2.5% glutaraldehyde, 2% paraformaldehyde, and 0.1 M sodium cacodylate buffer (pH 7.4, Electron Microscopy Sciences, Munich, Germany) overnight at 4°C. After rinsing in 0.1 M sodium cacodylate buffer, samples were post-fixed in 1% OsO4 for 1.5 hours at room temperature, washed in cacodylate buffer, and dehydrated with 50 % ethanol. Tissues were counterstained with 6% uranyl acetate dissolved in 70% ethanol (Serva, Heidelberg, Germany), followed by graded ethanol concentrations up to 100 % and Propylenoxide. The dehydrated samples were incubated in 2:1, 1:1, and 1:2 mixtures of propylene oxide and Epon resin (Serva) for 1 hour each. Finally, samples were infiltrated with pure Epon for 2 hours. Samples were embedded in fresh resin in block molds and cured for 3 days at 60°C. Semithin sections (500 nm) were cut on a Reichert Ultracut S (Leica Microsystems, Wetzlar, Germany) and stained with Richardson staining solution. Ultrathin sections (50 nm) were cut on the same Ultracut, collected on copper grids, and counterstained with Reynold’s lead citrate. Sections were analyzed with a Zeiss EM 900 transmission electron microscope (Zeiss) equipped with a 2k x 2k CCD camera.

### Statistical analyses

All data sets were tested for normal distribution (D’Agostino and Pearson test) and were subjected to outliers’ identification via ROUT method (Q=1%). *In vitro* assay data obtained from each HQ-treated group were normalized to the respective control untreated group for each cell line (represented as dotted line in the graphs and set as 1) to avoid artefacts deriving from different cell numbers,. The relative effects of HQ in each group were derived with paired Student T-test (#) compared to controls (dotted line). Differences between LR and HR groups were derived with unpaired Student T-test (*). Data showing JC-1 ratio, which is not dependent on number of cells, were analyzed with one-way ANOVA with Tukeýs multiple comparisons test. Relative mRNA expression was calculated with the ΔΔCt method, considering the average of all ΔCt of all LR RPE cells as controls. Gene expression changes were analyzed with one-way ANOVA with Tukeýs multiple comparisons test. Western Blot images were analyzed for signal quantification using Fiji (ImageJ), and relative protein expression levels were analyzed using one-way ANOVA with Tukeýs multiple comparisons test. Analyses were performed using GraphPad Prism 10 software. Data are shown as mean ± SEM. Individual data points represent biological replicates. Significance level was set as p ≤ 0.05.

## RESULTS

### iPSC-RPE carrying high genetic risk (HR) are more vulnerable to Hydroquinone mediated-damage

To assess the impact of genetic risk variants on RPE cells, we used iPSC-RPE lines and used the FH Y402H polymorphism as main criteria for classification into low risk (LR) or high risk (HR), where LR (n=3) are homozygous carriers of the FH 402Y and HR are homozygous carriers of FH 402H (n=5). Both groups present a mixed genotype at Chr10 for ARMS2 and donors were selected independently of their AMD status (detailed in Suppl. Figure 1A).

The RPE cells were evaluated for maturity based on established criteria. Both LR and HR RPE cells exhibited mature characteristics, including pigmentation (Figure 1A-D), hexagonal shape confirmed by immunostaining of tight-junction protein ZO-1 (Figure 1E-H), and monolayer formation with normal apical microvilli (Figure 1I-J). Strong tight junctions and proper polarization were confirmed by proper trans-epithelial resistance (Figure 1L), consistent with previous studies. In addition, the cell lines also showed a polarized secretion of PEDF, a characteristic of mature RPE cells, with higher levels in the apical extracellular compartment (Figure 1K). RPE markers were also validated by gene expression analyses (Suppl. Figure 1B). Given the hypothesized role of FH in RPE pathology during AMD progression, we confirmed that RPE cells do express *CFH* mRNA (Suppl. Figure 1C), exhibit FH in protein lysates from both LR and HR iPSC-RPE cells (Figure 1M) and secretion of C3 (Suppl. Figure 1D).

**Figure 1.**
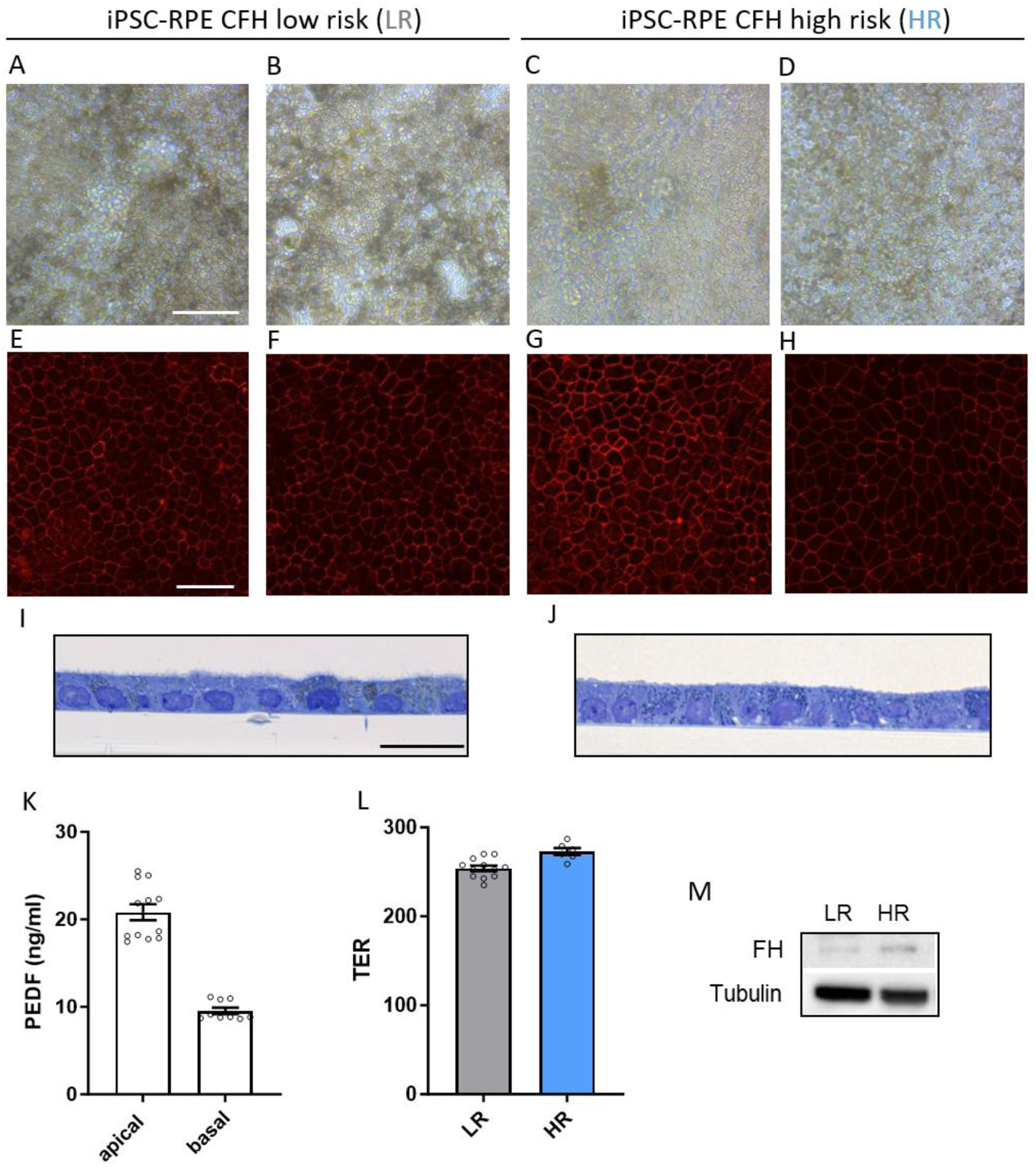
Generation of CFH low risk (LR) and high risk (HR) iPSC-RPE. **A-D** Representative light microscopy images of iPSC-RPE cells derived from LR donors iPSC-RPE2 (A), iPSC-PRE3 (B) and HR donors iPSC-RPE5 (C), iPSC-PRE8 (D). Images show proper RPE cuboidal shape and visible pigmentation. Scale bar = 100 µM. **E-H** Representative immunostaining images of RPE marker ZO-1 in LR donors iPSC-RPE1 (E), iPSC-PRE3 (F), and HR donors iPSC-RPE6 (G), iPSC-PRE7 (H). Scale bar = 50 µM. **I-J** Transmission electron micrographs semithin sections of LR donors iPSC-RPE3 (I) and HR donors iPSC-RPE8 (J). Images show an RPE monolayer and microvilli typical of RPE cells. Scale bar = 20 µM. **K** Pigment epithelium-derived factor (PEDF) ELISA of apical and basal supernatant. Data collected from iPSC-RPE3, 5, 6, 7, 8. **L** Trans-epithelial resistance (TER) measured in iPSC-RPE3, 5, 8. **M** Representative WB images of FH levels in iPSC-RPE1 and iPSC-RPE5 cellular lysates. Tubulin was used as housekeeping control.

Our objective was to investigate how genetic predisposition influences AMD pathology at the RPE level. We hypothesized that RPE cells with high genetic risk (HR) may be more susceptible to damage from adverse conditions such as aging or unhealthy lifestyle factors. One of the key features of these AMD risk factors is increased oxidative stress. As such, we combined genetic risk with HQ, an oxidative agent found in cigarette smoke: smoking is itself a large contributor of AMD-risk [31]. We treated LR and HR iPSC-RPE cells with increasing concentrations of HQ (100 and 200 µM) for 48 hours and assessed the RPE stress response (Figure 2A). Initially, we compared membrane damage levels in a cytotoxicity assay by comparing HQ-treated LR and HQ-treated HR iPSC-RPE cells and their respective non HQ-treated controls (dotted line set as 1). Only HR iPSC-RPE cells exhibited cytotoxicity at 200 µM HQ (Figure 2B). As apoptosis is a primary cell death response to oxidative stress, we examined apoptosis markers in both LR and HR iPSC-RPE cells. At the lower HQ concentration (100 µM), both LR and HR cells showed significant caspase 3/7 activation, with a more pronounced response in HR iPSC-RPE cells. At the higher concentration (200 µM), only HR iPSC-RPE cells exhibited significantly elevated caspase levels compared to untreated HR controls (Figure 2C). Additionally, we investigated the protein expression levels of Bax (pro-apoptotic) and Bcl-2 (anti-apoptotic). We calculated the Bax/Bcl-2 ratio, which was elevated in HR iPSC-RPE cells towards Bax expression even under untreated conditions, indicating a predisposition to higher apoptotic levels. This ratio remained unchanged in HQ-treated LR iPSC-RPE cells but increased significantly in HR iPSC-RPE cells after 200 µM HQ exposure (Figure 2D-E).

**Figure 2.**
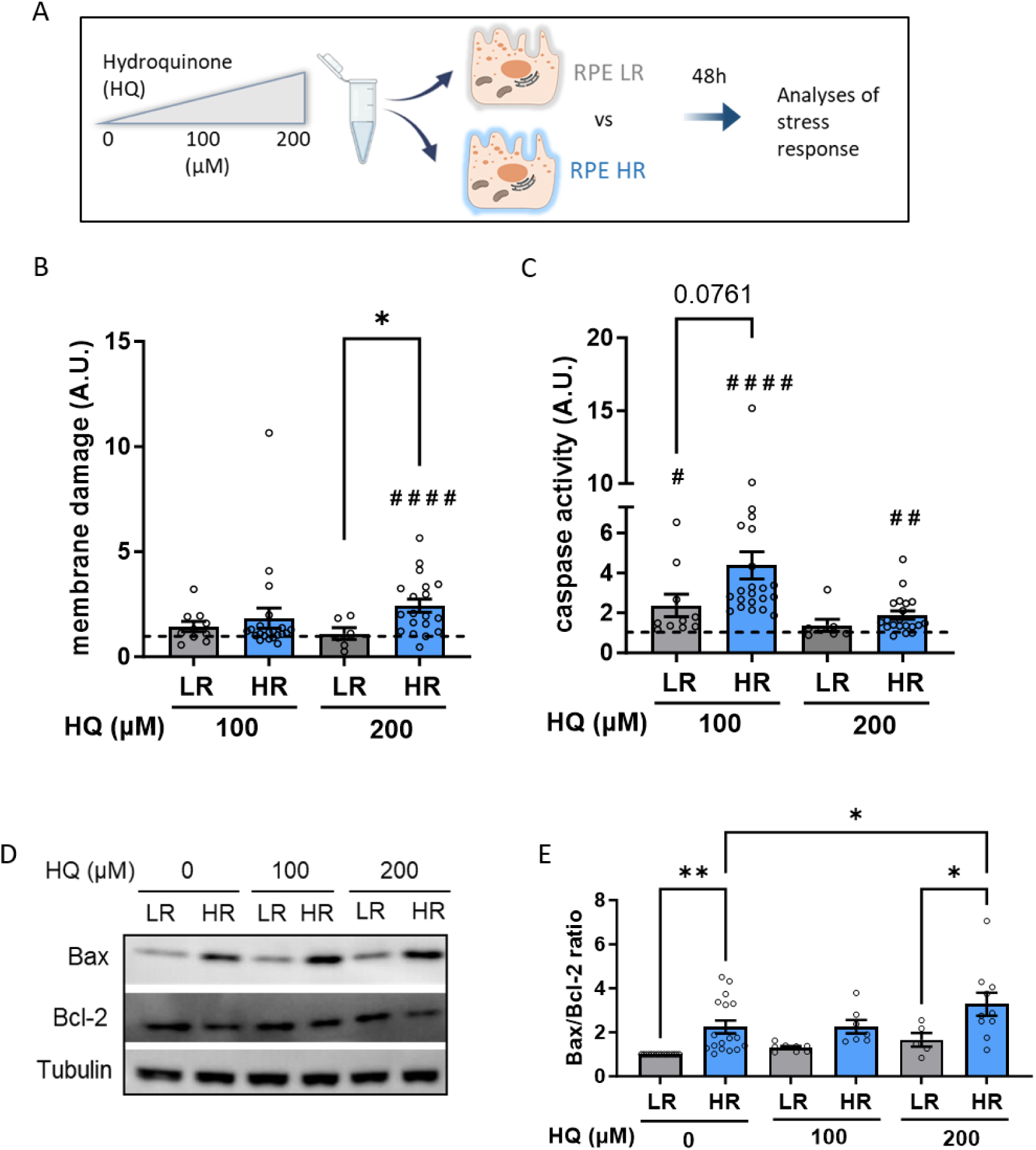
Hydroquinone (HQ) induces cellular damage and apoptosis in CFH high risk (HR) iPSC-RPE. **A** Schematic representation of the experimental set up. **B-C** Membrane damage assessed by cytotoxicity assay GF-AFC (B) and Caspase-3 activity (C) in LR and HR iPSC-RPE. Data points were collected from LR iPSC-RPE1, 2, 3 (n=3 biological replicates) and from HR iPSC-RPE4, 5, 6, 7, 8 (n=5 biological replicates). HQ-treated relative values are normalized to the respective untreated controls (dotted line) in each individual experiment for each cell line. HQ effects in each group were assessed with paired Student’s t-test (#) compared to controls (dotted line). Differences between LR and HR groups were determined with unpaired Student’s t-test (*). **D-E** Representative WB images (D) of Bax and Bcl-2 levels LR and HR iPSC-RPE cells treated with HQ. Data points were collected from LR iPSC-RPE1, 2, 3 (n=3 biological replicates) and from HR iPSC-RPE4, 5, 6, 7, 8 (n=5 biological replicates). Tubulin was used as housekeeping control. Quantification of Bax/Bcl-2 ratio is shown in E. Differences between LR and HR groups were assessed with one-way ANOVA (*). Data are shown as mean ± SEM.

### iPSC-RPE carrying high genetic risk (HR) show mitochondrial impairment

The ability to maintain metabolic balance even when exposed to oxidative conditions and an adverse environment is crucial for RPE homeostasis. This is the cellular environment during AMD progression. Healthy RPE cells are supposed to cope with stress and minimally vary their metabolic functions. To assess whether the susceptibility to HQ in HR iPSC-RPE cells was reflected at a metabolic level, we used *in vitro* assays to measure ATP levels and mitochondrial polarization. Indeed, we observed that, although both LR and HR cells exhibited reduced ATP levels after HQ exposure, the reduction was more pronounced in HR cells (Figure 3A). To investigate mitochondrial integrity, we used a JC1 assay, which allows the assessment of mitochondrial membrane potential, and we demonstrated that only the HR cells, when treated with higher concentrations of HQ, showed a decreasing trend in mitochondrial membrane depolarization (Figure 3B). Interestingly, we observed mitochondrial impairment in HR cells even under control conditions, which may predispose these cells to greater damage in response to oxidative insults such as HQ exposure. Using electron microscopy (EM), we observed that mitochondria in LR and HR RPE were localized differentially. Classically, mitochondria are localised in the basal compartments of RPE cells, as was indeed observed in LR RPE cells, whereas HR iPSC-PRE showed a partial localisation of mitochondria in the apical compartment (Figs. 3C-D, mitochondria highlighted in green). Furthermore, the mitochondria are altered in HR RPE cells, as previously shown [15]. Mitochondria in LR RPE cells appear to be organised, with a double membrane and clear cristae, irrespective of basal or apical localisation (Figures E-F, mitochondria marked with white arrowheads). On the other hand, morphology of mitochondria in HR RPE cells appears polymorphic, varying in shape and size (Figure 3G-H, polymorphic mitochondria marked with white arrowheads). Mitochondria in HR RPE cells have vacuoles, indicating swollen mitochondria; the inner cristae structure is damaged (Figure 3G-H, swollen mitochondria marked with black arrowheads). Another feature in HR RPE cells was the presence of bent mitochondria (Figure 3G-H, bent mitochondria marked with asterisk), which has been previously observed in photoreceptor mitochondria of older mice and is associated with mitochondrial motility [32]. Moreover, in both groups it was possible to detect pinched mitochondria, which are an indication of fission and fusion, and we found those features were present in higher number in the HR RPE cells, indicating altered mitochondrial dynamics. Therefore, we comparably investigated a possible involvement of mitophagy in the differential response to stress. Although no major differences in the level of the main mitophagy markers PINK and PARKIN [33] were observed in the untreated condition, LR and HR RPE cells differentially regulated the expression of those genes in response to HQ. In LR cells we found PINK1 downregulated while it remained unchanged in HR cells. HR downregulated PARKIN in response to HQ, while LR did not. In parallel, the levels of PGC1a, a transcription factor regulating mitochondrial biogenesis [34] was reduced in both LR and HR after HQ treatment (Figure 3I-K).

**Figure 3.**
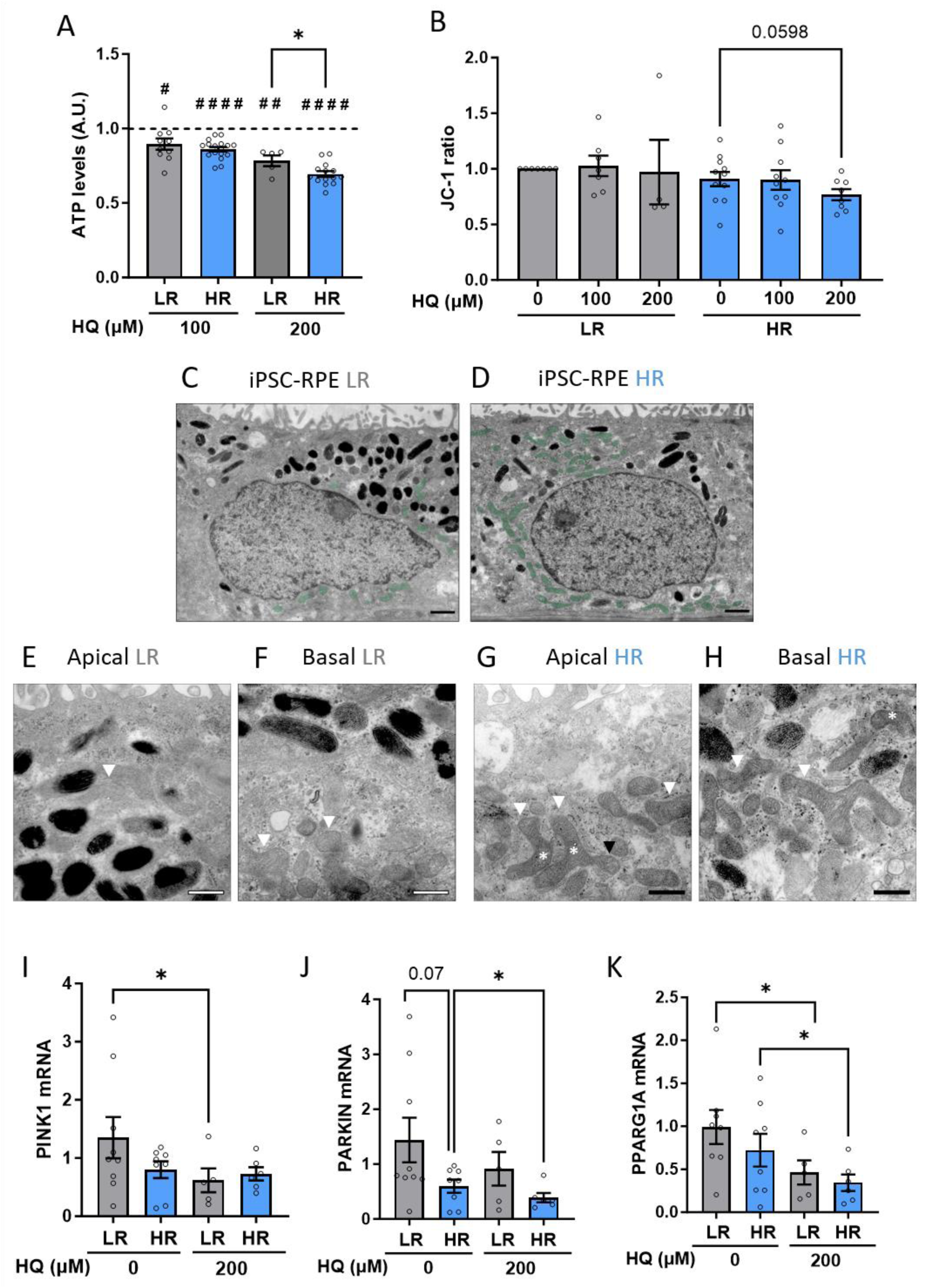
CFH high risk (HR) iPSC-RPE cells show impaired mitochondria homeostasis. **A** ATP levels assessed by cell titer assay in LR and HR iPSC-RPE. HQ-treated relative values are normalized to the respective untreated controls (dotted line) in each individual experiment for each cell line. HQ effects in each group were determined with paired Student’s t-test (#) compared to controls (dotted line). Differences between LR and HR groups were determined with unpaired Student T-test (*). **B** JC-1 fluorescence measurements were recorded, and ratio was determined. HQ effects in each group were assessed with paired Student’s t-test (#) compared to untreated controls. **C-H** Representative EM images of LR iPSC-RPE3 and HR iPSC-RPE8 showing mitochondria abnormalities. C-D show the mitochondria localization (highlighted in green) in the apical compartment in HR (D) compared to basal localization (C). scale bar = 1µm. E-H show mitochondria morphology. White arrowheads in E-F mark healthy mitochondria in LR iPSC-RPE cells. In G-H white arrowheads mark polymorphic mitochondria, black arrowheads mark swollen mitochondria, asterisks mark bent mitochondria. Scale bar = 500nm. **I-J** Gene expression levels of PINK (I), PARKIN (K) and PPARG1A (J) analyzed via RT-qPCR in LR and HR iPSC-RPE treated with HQ. Significance was determined with one-way ANOVA (*). Data are shown as mean ± SEM. Data points were collected from LR iPSC-RPE1, 2, 3 (n=3 biological replicates) and from HR iPSC-RPE4, 5, 6, 7, 8 (n=5 biological replicates).

### HQ-mediated damage in iPSC-RPE carrying high genetic risk (HR) is not dependent on extracellular FH or complement sources

Recent research has proven the existence of non-canonical functions of FH, which are distinct from its well-established role as a complement system inhibitor in the extracellular space. These non-canonical functions suggest that FH has a role in influencing cellular homeostasis within the intracellular environment. Our study investigated whether the increased susceptibility of HR iPSC-RPE cells to Hydroquinone (HQ)-induced damage could be linked to these non-canonical functions of FH.

We first exposed HR iPSC-RPE cells to HQ in the presence of plasma-purified soluble FH to determine if FH could attenuate the HR phenotype (Figure 4A). Notably, HR iPSC-RPE cells exposed to purified FH exhibited FH in their cellular lysates, indicating either cellular uptake or binding to the cellular membrane (Figure 4B). Purified FH appeared to have a higher molecular weight, likely due to differential glycosylation, as observed and hypothesized in other models [35, 36]. However, no rescue effect on cytotoxicity or caspase activation was observed in HR iPSC-RPE cells exposed to HQ and purified FH (Figure 4C-D).

**Figure 4.**
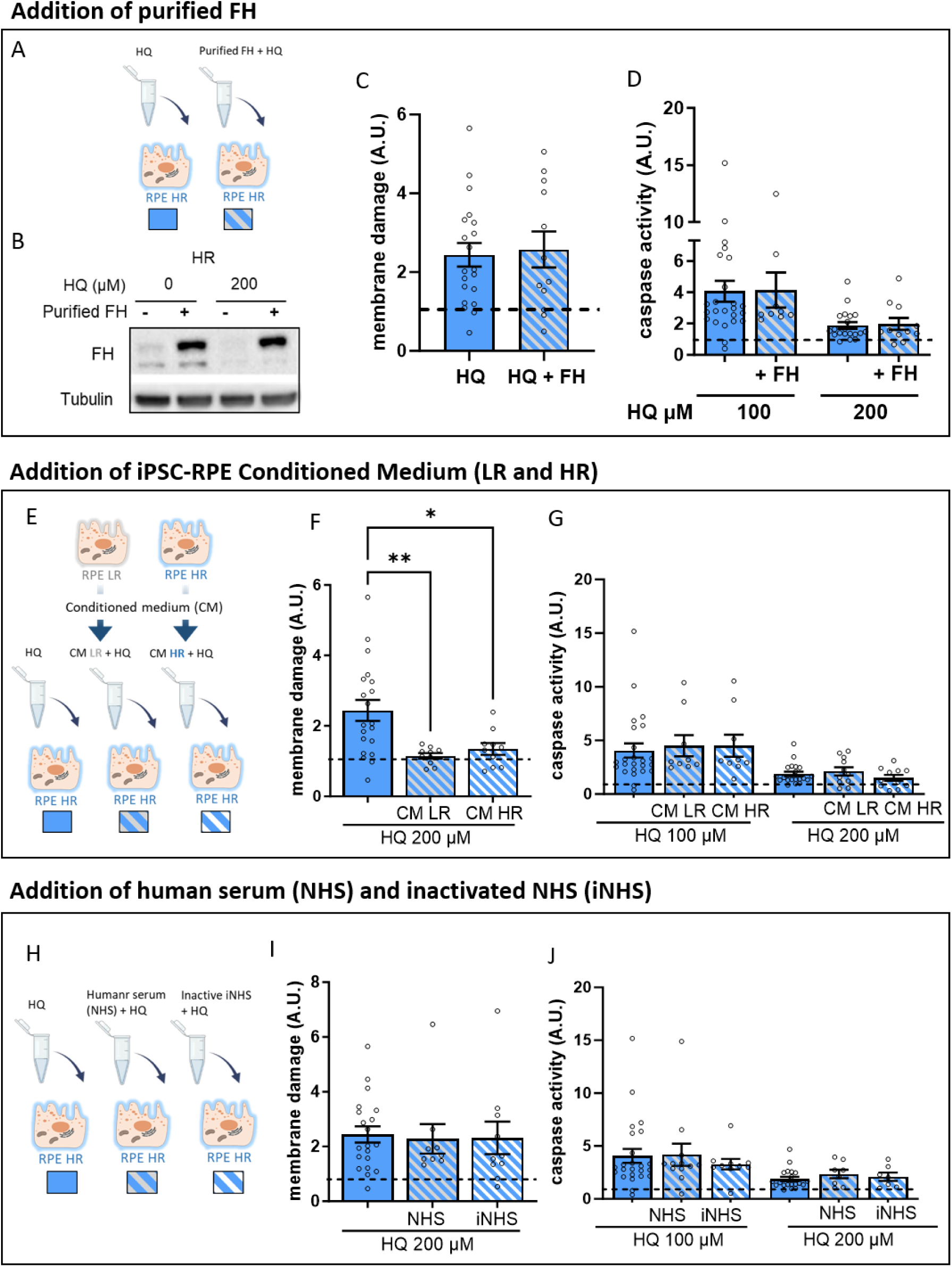
Addition of extracellular FH and complement sources does not affect the vulnerability to HQ-mediated stress in HR iPCS-RPE. **A** Schematic representation of the experimental set up of FH supplementation. **B-C** Membrane damage assessed by cytotoxicity assay GF-AFC (B) and Caspase-3 activity (C) in HQ-treated HR iPSC-RPE with supplementation of purified FH. Values are normalized to the respective controls (dotted line) in each individual experiment for each cell line. Data are shown as mean ± SEM. **D** Schematic representation of the LR and HR iPSC-RPE conditioned medium supplementation experimental set up. **E-F** Membrane damage assessed by cytotoxicity assay GF-AFC (B) and caspase-3 activity (C) in HQ-treated HR iPSC-RPE with supplementation of LR and HR conditioned medium. Values are normalized to the respective controls (dotted line) in each individual experiment for each cell line. Significance was derived with 2-way ANOVA. **H** Schematic representation of the experimental set up of HQ-treated HR iPSC-RPE supplemented with NHS and iNHS. **I-J** Membrane damage assessed by cytotoxicity assay GF-AFC (B) and caspase-3 activity (C) in HQ-treated HR iPSC-RPE with supplementation of LR and HR conditioned medium. Values are normalized to the respective controls (dotted line) in each individual experiment for each cell line. Data points were collected from HR iPSC-RPE5, 6, 7, 8 (n=4 biological replicates). Data are shown as mean ± SEM.

We then treated HR iPSC-RPE cells with HQ in combination with conditioned medium collected from either LR or HR iPSC-RPE cells (Figure 4E). The use of conditioned medium provided mixed results, with reduced cytotoxicity in HR iPSC-RPE cells exposed to both LR and HR conditioned media (Figure 4F), yet caspase activation remained elevated in HR iPSC-RPE treated with HQ (Figure 4G). This suggests that neither secreted FH variant is able to rescue our observed HQ-induced elevation of caspase activity; rather, the susceptibility appears to be intrinsically linked to the endogenous expression of the HR variant in iPSC-RPE cells.

Given FH’s canonical role as a complement inhibitor, we assessed whether complement activation contributes to the observed damage in HR iPSC-RPE cells. We performed HQ treatment experiments in the presence of normal human serum (NHS), in which complement activation is possible, and inactivated NHS (iNHS), in which complement activation is limited (Figure 4H). Our findings indicate that increased cytotoxicity and caspase activation in HQ-exposed HR iPSC-RPE cells are independent of complement system activation, as no differences were observed between treatments with NHS or iNHS (Figure 4I-J).

### Autophagy flux is impaired in iPSC-RPE carrying high genetic risk (HR)

To further elucidate the mechanisms underlying the increased vulnerability of HR iPSC-RPE cells to hydroquinone (HQ)-induced stress, we investigated the oxidative stress response. Given that HQ is an oxidative agent, we first assessed oxidative stress by measuring hydrogen peroxide (H₂O₂) levels. As expected, H₂O₂ levels increased in both LR and HR iPSC-RPE cells in a concentration-dependent manner; however, no significant differences were found between LR and HR cells (Suppl. Figure 2A). We also analyzed the expression of key antioxidant components, including catalase (CAT, Suppl. Figure 2B), glutathione peroxidase 1 (GPX1, Suppl. Figure 2C), and superoxide dismutase 2 (SOD2, Suppl. Figure 2D-E). CAT and GPX1 gene expression levels were similar between LR and HR iPSC-RPE cells. However, in both the untreated and HQ-treated conditions, a modest but significant reduction in SOD2 protein levels was observed in HR iPSC-RPE cells compared to LR cells.

Under conditions of oxidative stress, the endoplasmic reticulum (ER), which is crucial for protein synthesis, folding, and regulation, may become dysfunctional under oxidative stress conditions. To explore this possibility, we assessed the ER stress response and the unfolded protein response (UPR) by examining the expression of the chaperone Binding immunoglobulin protein (BiP), an initial sensor of unfolded proteins, and inositol-requiring enzyme 1 α (IRE1α), a major UPR effector (Suppl. Figure 3A-C). Although there was a significant increase in BiP in LR cells after treatment with HQ, no significant differences were noted between HR and LR iPSC-RPE cells. To ensure that HR iPSC-RPE cells do not have an inherently impaired ER stress response, we treated both cell groups with Tunicamycin (TM), a known ER stress inducer. HR iPSC-RPE cells showed a small, but significant elevated cytotoxicity and both LR and HR iPSC-RPE cells showed caspase activation in response to TM (Suppl. Figure 3D-E). The Bax/Bcl-2 ratio was, however, higher in TM-treated HR iPSC-RPE cells compared to TM-treated LR cells, reflecting the differences observed in basal conditions (Suppl. Figure 3F-G). Both cell groups demonstrated a similar UPR activation in response to TM, with significant upregulation of BiP and IRE1α (Suppl. Figure 3 H-J).

Lastly, we examined the impact of oxidative/proteotoxic stress on the autophagic flux. Both LR and HR iPSC-RPE cells exhibited accumulation of LC3-II and p62 in response to HQ treatment, indicating increased autophagy (Figure 5A-C). However, detailed analysis using Bafilomycin A1 (BafA1), a commonly used autophagy inhibitor, revealed that LR iPSC-RPE cells had a higher capacity for autophagy compared to HR iPSC-RPE cells, as evidenced by a significant reduction in LC3-II levels in BafA1-treated HR iPSC-RPE cells (Figure 5D-E).

**Figure 5.**
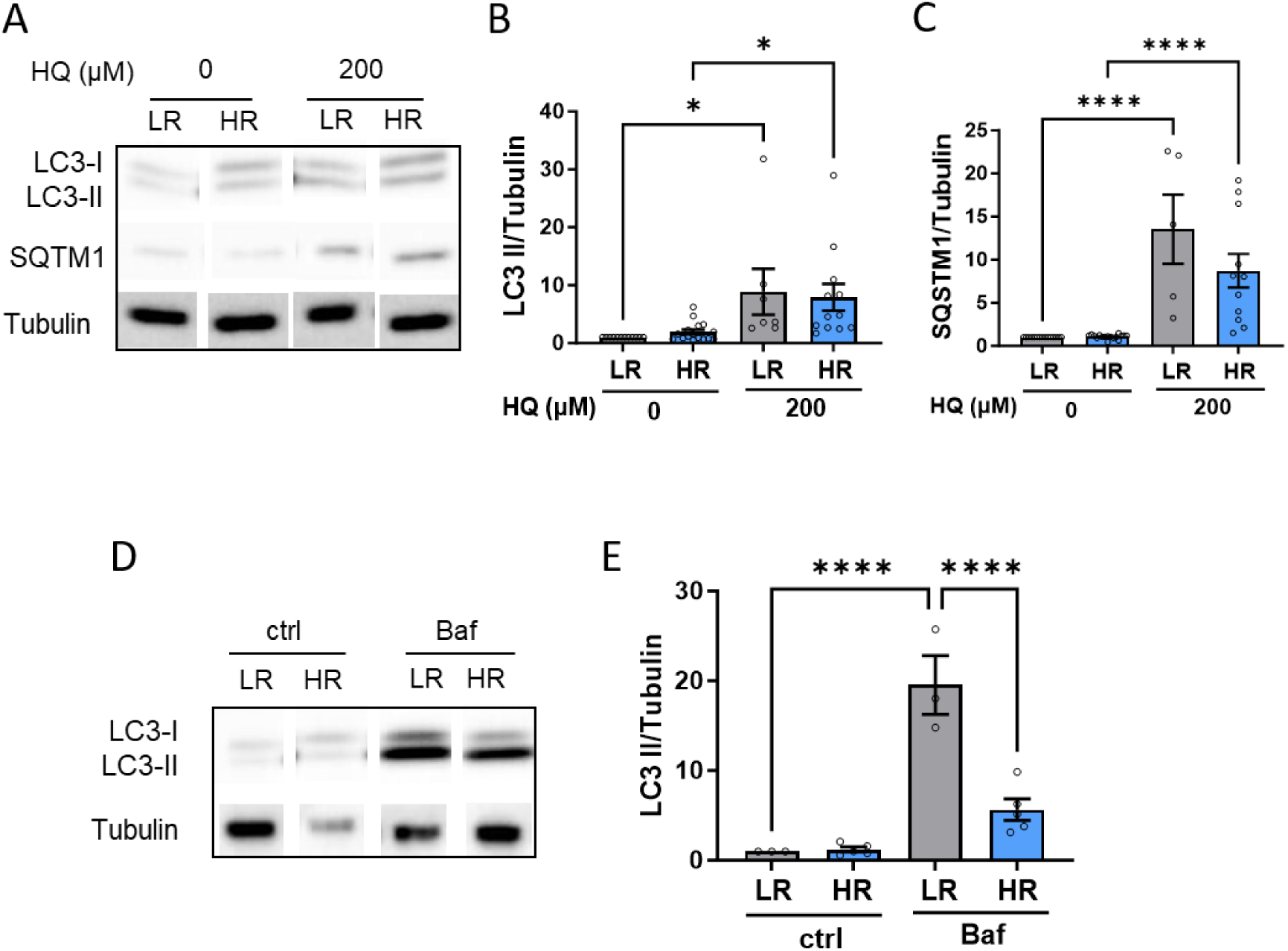
HR iPSC-RPE shows impaired autophagic flux. **A-C** Representative WB images (A) of LC3 and SQSTM1 in LR and HR iPSC-RPE cells treated with HQ. Tubulin was used as housekeeping. Quantification is shown for LC3 (B) and SQSTM1 (C). Data points were collected from LR iPSC-RPE1, 2, 3 (n=3 biological replicates) and from HR iPSC-RPE4, 5, 6, 7, 8 (n=5 biological replicates). **D-E** Representative WB images (D) of LC3 LR and HR iPSC-RPE cells treated with Bafilomycin. Tubulin was used as housekeeping. Quantification are shown for LC3 (E). Data points were collected from LR iPSC-RPE1, 3 (n=2 biological replicates) and from HR iPSC-RPE6, 8 (n=2 biological replicates). Differences between groups were determined with one-way ANOVA (*). Data are shown as mean ± SEM.

## Discussion

AMD leads to a devastating loss of visual acuity and central vision, as a result of a complex combination of risk factors, including genetic predisposition, age and unhealthy life-style habits [37, 38]. Smoking has been identified as a major risk that synergizes with genetic risks accelerating the onset and progression of the disease [37]. However, it remains enigmatic so far as to how, where and when the combination of genetic and life-style-associated risk contributes to the disease. About 50% of AMD patients present polymorphisms in the complement factor H (*CFH*) gene, an inhibitory component of the complement pathway. Risk variants in the *CFH* gene lead to reduced inhibitory function of the Factor H protein (FH) [7]. Until recently, FH risk variants were thought to confer susceptibility to AMD solely through over-activation of the complement system, and clinical trials were designed to inhibit this process.

Two recent studies we could show that local loss of *CFH* in RPE results in pathophysiological consequences not linked to the complement pathway. Loss of *CFH* in RPE cells results in an increase of inflammatory cytokines and chemokines not directly linked to the complement pathway [39]. Searching for the causative molecular mechanisms, we identified the NF-ƙB pathway as the major pathway involved in those changes. We could further show that *CFH* loss in RPE cells has severe consequences on energy metabolism, resulting in a reduction of both glycolysis and mitochondria respiration [23]. Using hTERT-RPE1 cells we could show, that loss of FH expression in these cells results in a higher vulnerability to H_2_O_2_ and an impairment in mitochondrial metabolism [23]. These studies, however, were performed by *CFH* knockout within RPE cells. A variety of studies with ARPE19 cells, found a higher cell damage in response to a multiplicity of oxidative stimuli (H_2_O_2_, tBH, 4-HNE, HQ, CSE) [26, 40–42].

To get closer to human pathology, we employed RPE cells derived from induced pluripotent stem cells (iPSC) generated from donors, carrying either homozygous 402Y (low risk) or 402H (high risk) variant of the *CFH* gene. RPE cells derived thereof were treated with Hydroquinone (HQ), a component of cigarette smoke, to induce oxidative damage. RPE cells carrying the high genetic risk in *CFH* proved more vulnerable to oxidative insult when exposed to HQ, as demonstrated by increased cytotoxicity and caspase activation, compared to the low risk RPE cells. Considering the known connection of oxidative stress to proteotoxic stress and degrading processes, we investigated the unfolded protein response (UPR) and autophagy. When exposed to HQ, RPE cells showed an increase in autophagy markers, however HR RPE cells showed an overall reduced autophagic flux. The degree of cellular susceptibility to oxidative stress turned out to be solely conferred by cell-endogenously produced intracellular FH, as soluble FH protein and other complement sources did not influence the response to HQ. Moreover, having both, high-risk and low-risk genotypes for Chr10q26 (*ARMS2/Htra1*) included, we can show, that the degree of cellular susceptibility to oxidative stress solely associates to the risk haplotype on chromosome Chr1q32, but not to chromosome 10q26.

The risk loci on Chr1q32 includes the protective I62V polymorphism and a risk Y402H polymorphism in FH, and a FHR deletion region, all of which contribute to the genetic haplotype with the highest association with AMD [43]. However, most of the research to date has focused on the FH Y402H polymorphism, and properly genotyped iPSC-RPE with fully defined haplotypes would be needed to fully elucidate the contribution of each of the individual risk polymorphisms on Chr1q32 to RPE homeostasis. It has been shown that FH Y402H leads to excessive activation of the alternative pathway of the complement system and products of complement activation may accumulate at a systemic level or at the Bruch’s-choroid interface, or locally around the RPE cells themselves [44]. However, knowledge gained in recent years has revealed novel non-canonical functions of the complement system [13, 45]. These non-canonical cellular functions include T-cell activation and metabolism [46] and regulation of differential transcriptional activity in cancer cells [47]. Autophagy and lysosomal capacity have been found to be controlled intracellularly by C3 in pancreatic β-cells [48]. These are only some of the recent discoveries that point out a prominent role of the complosome (active intracellular complement components) in tissue homeostasis and disease pathology [45]. Some of those functions have been found to depend on intracellular complement system activation, while others simply depend on the intracellular activity of complement proteins, exerting non-canonical functions.

In our study, HR RPE cells showed increased cytotoxicity in response to HQ in a dose-dependent manner. In response to HQ, we observed an increase in H_2_O_2_ in both groups of cells (LR and HR), confirming that the degree of cell damage is not determined by the levels of ROS per se, but rather by the differences in the oxidative stress response. HQ-induced cell damage by mediating apoptosis was confirmed by caspase-3 activation and an increase in the Bax-Bcl2 ratio in HR RPE cells. The addition of exogenous FH or conditioned medium containing FH did not rescue either cytotoxicity or caspase activation in a risk variant-dependent manner. These results are consistent with previous observations in colorectal cancer line - ccCRC cells, endothelial cells, and immortalized fibroblasts [47, 49]. In these cell types, reduced levels of intracellular FH were associated with reduced viability whereas the addition of FH had no effect [47]. On the contrary, exogenous FH was able to reverse the H_2_O_2_ mediated damage in ARPE19 cells and the 4-HNE damage in iPSC-RPE cells [40, 50] most likely by reducing the pro-inflammatory status in response to the oxidative stimuli. However, in our study, no differences in inflammation were observed when comparing iPSC-RPE LR and HR, either under physiological conditions or when stressed with HQ (data not shown). This was confirmed in a parallel study where IL-6 levels were similar when comparing iPSC-RPE cells derived from donors with different disease states and genotypes [14].

iPSC-RPE cells and primary RPE cells carrying the Y402H polymorphism were shown to accumulate swollen lysosomes, show reduced lysosomal activity, damaged mitochondria and impaired autophagy: all changes that affect the metabolic balance between the RPE and the neuroretina [14–16, 51]. However, whether these features are regulated by complement inside the RPE cells or mediated by complement dysregulation outside the cells remained to be clarified in these studies. Consistent with these findings, we also observed mitochondrial abnormalities and reduced autophagic flux in HR iPSC-RPE cells. In particular, we observed reduced autophagic flux and imbalanced mitochondrial dynamics in HR RPE cells, which may be the manifestation of reduced autophagic and lysosomal function and the cause of mitochondrial damage, as previously suggested [51]. Taken together, these studies point to a severe physiological imbalance of RPE cells carrying a risk haplotype on Chr1q32 in response to exogenous lifestyle-related stressors.

Our data suggest that endogenously expressed FH protects the RPE. It remains to be shown whether and how this protection of the RPE contributes to protect the neuroretina. Recent work has shown that a dysfunctional RPE triggers photoreceptor degeneration and, in particular, rod cell loss in the parafovea of AMD patients [52]. In this study using a complement inhibitor, Pegcetacoplan directed against C3 conversion, drug-treated eyes showed a significantly slower GA lesion progression rate compared with sham-treated controls. To date, inhibition of the complement system has been the main therapeutic target for dry AMD. However, its efficacy may be highly dependent on the patient’s genotype and stage of the disease. Individuals with AMD may be responsive to complement inhibition in the early stages of the disease, when the influence of life-style factors and aging is less. Consequently the susceptibility to stress due to the intracellular FH Y402H variant has a limited effect. This is supported by the fact that both LR and HR iPSC-RPE cells were damaged by the extracellular complement system in the absence of additional stress [16]. As the disease progresses or in the presence of combined higher risk, the effect of intracellular FH on RPE physiology, which is most likely independent of complement activation, may no longer respond to complement inhibitory therapies. This hypothesis reinforces the need to consider different avenues for AMD treatment, such as patient stratification or combination therapy, combining complement inhibition with approaches that protect RPE physiology.

---

**Supplementary Figure 1.**
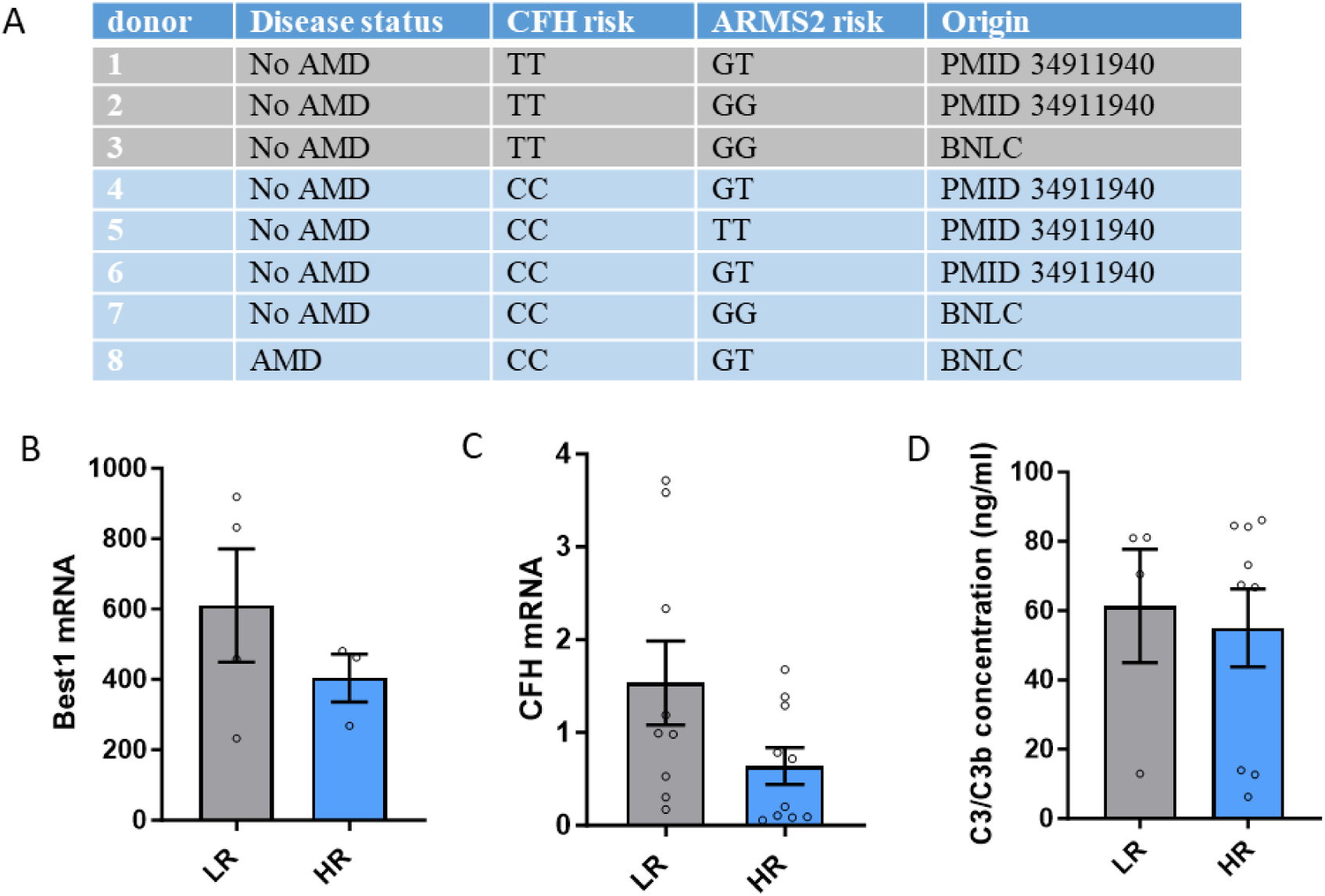
Generation of iPSC-RPE cells. **A** Summary table with the information relative to genetics and origin of the iPSC lines. Spanish National Stem Cell Bank (BNLC). **B** Gene expression levels of RPE marker BEST1 analyzed via RT-qPCR in LR and HR iPSC-RPE. Data are normalized to Best1 expression in hTERT-RPE1 cells. **C** Gene expression levels of CFH analyzed via RT-qPCR in LR and HR iPSC-RPE. **D** C3/C3b levels analyzed by ELISA in cell culture supernatants in LR and HR iPSC-RPE. Data are shown as mean ± SEM.

**Supplementary Figure 2.**
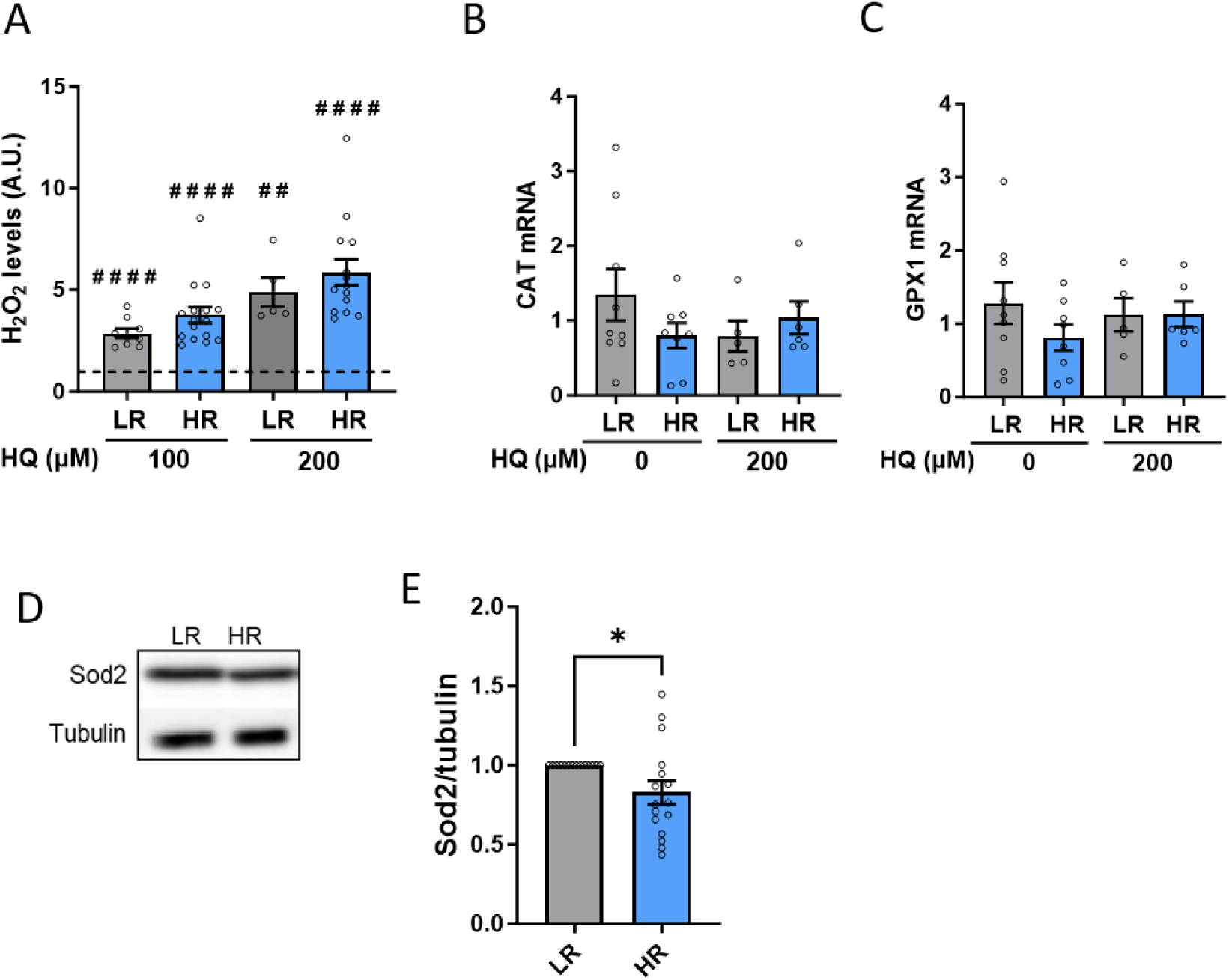
Hydroquinone (HQ) impact on oxidative stress in LR and HR iPSC-RPE. **A** H2O2 levels assay in LR and HR iPSC-RPE. HQ-treated relative values are normalized to the respective controls (dotted line) in each individual experiment for each cell line. HQ effects in each group were assessed with paired Student’s t-test (#) compared to controls (dotted line). **B-C** Gene expression levels of CAT (B) and GPX1 (C) analyzed via RT-qPCR in LR and HR iPSC-RPE treated with HQ. **D-E** Representative WB images (D) of Sod2 levels LR and HR iPSC-RPE. Tubulin was used as housekeeping control. Quantification of Sod2 levels is shown in E. Differences between LR and HR groups were determined with one-way ANOVA (*). Data points were collected from LR iPSC-RPE1, 2, 3 (n=3 biological replicates) and from HR iPSC-RPE4, 5, 6, 7, 8 (n=5 biological replicates). Data are shown as mean ± SEM.

**Supplementary Figure 3.**
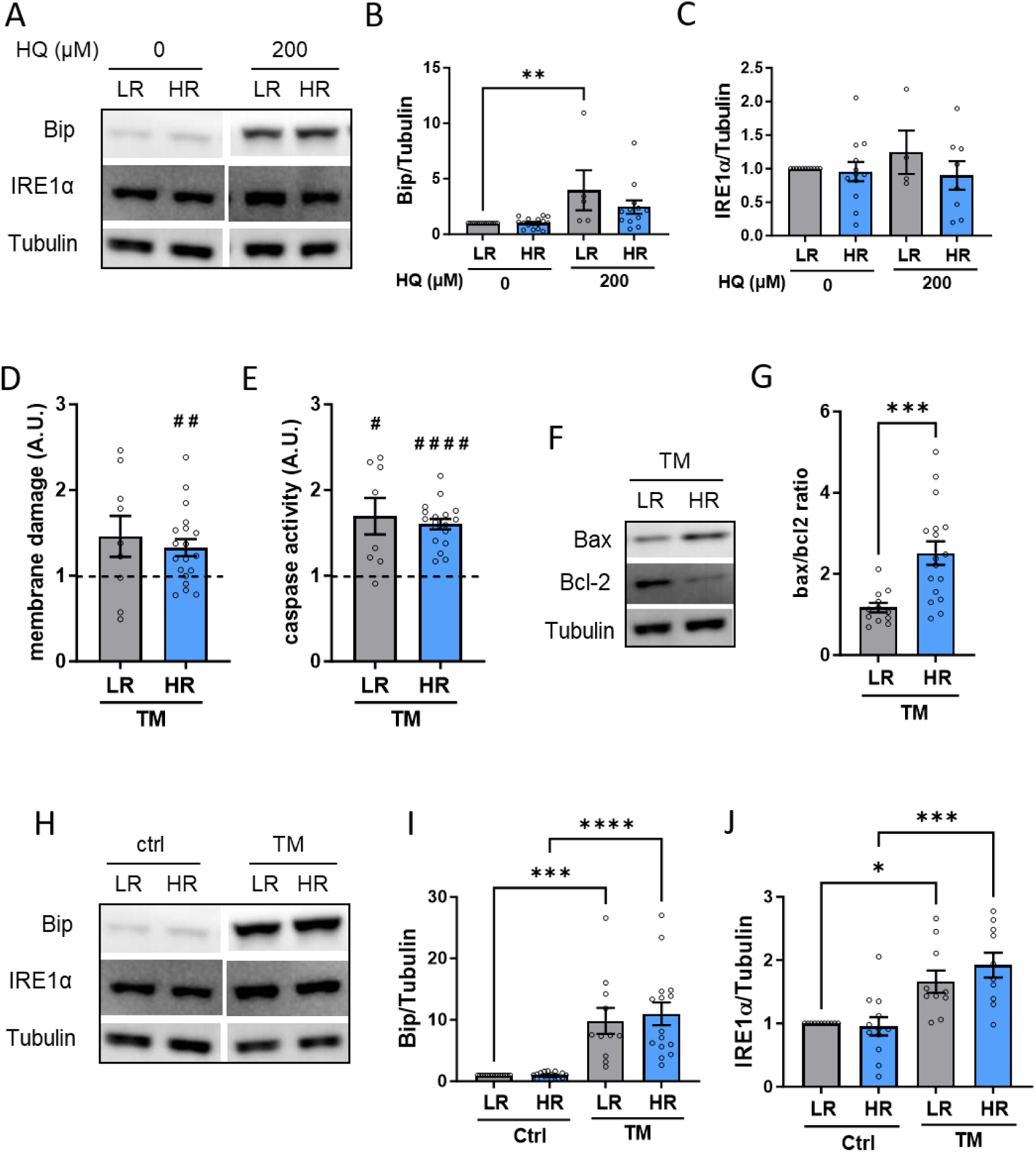
Impact of HQ and TM on the UPR response of LR and HR iPSC-RPE. **A-C** Representative WB images (A) of Bip and IRE1α levels in LR and HR iPSC-RPE cells treated with HQ. Tubulin was used as housekeeping control. Quantification of Bip (B) and IRE1α (C) levels are shown. Differences between LR and HR groups were assessed with one-way ANOVA (*). Data are shown as mean ± SEM. **D-E** Membrane damage assessed by cytotoxicity assay GF-AFC (D) and caspase3 activity (E). TM-treated relative values are normalized to the respective controls (dotted line) in each individual experiment for each cell line. TM effects in each group were determined with paired Student’s t-test (#) compared to controls (dotted line). **H-J** Representative WB images (H) of Bip and IRE1α levels in LR and HR iPSC-RPE cells treated with TM. Tubulin was used as housekeeping control. Quantification of Bip (I) and IRE1α (J) levels are shown. Differences between LR and HR groups were assessed with one-way ANOVA (*). Data are shown as mean ± SEM. Data points were collected from LR iPSC-RPE1, 2, 3 (n=3 biological replicates) and from HR iPSC-RPE4, 5, 6, 7, 8 (n=5 biological replicates).

